# Phylogeography and genetic structure of the steppe marmot (*Marmota bobak* Müller, 1776)

**DOI:** 10.1101/2024.12.12.628213

**Authors:** Oleg Brandler, Valentina Tambovtseva, Andrey Tukhbatullin, Vadim Rumyantsev, Alexey Grachov, Svetlana Kapustina

## Abstract

The steppe marmot *Marmota bobak* is a key species in the steppe ecosystems of Eurasia and an important object of economic and conservation practice. Its wide distribution is separated by many ecological and geographical barriers. The studied morphological variability is characterized by clinal variation of characters and indistinct differences. The subspecies system, including three subspecies, has no clear spatial boundaries and is controversial. The genetic variability of *M. bobak* has not been extensively studied. We first investigated the genetic variability of *M. bobak* throughout its range using three traditional molecular markers of the mitochondrial genome *COI, cytb*, and a control region of mtDNA. We found two phylogroups in the genetic structure of this species, which are geographically arranged in an unexpected way. Also, the character of intraspecific genetic variability does not agree with the accepted subspecies division; individuals from terra typica of all three described subspecies belong to the same phyletic lineage. Further studies are needed to develop the subspecies system of *M. bobak*.

## INTRODUCTION

Marmots (genus *Marmota*) are the largest representatives of ground squirrels (tribe Marmotini, subfamily Xerinae), widely distributed in northern Eurasia and North America (Thorington et al., 2012). The bobak group, including the steppe marmot or baibak (*M. bobak* Müll. 1776), gray marmot (*M. baibacina* Kastschenko 1899), and forest-steppe marmot (*M. kastschenkoi* Stroganov et Yudin 1956), is the most evolutionarily young among Palaearctic marmots (Mills et al., 2023). This is expressed in insufficiently clear morphological differences and the presence of transitional forms (Bibikow, 1996).

Steppe marmot is adapted to inhabit open steppe plains or weakly hilly landscapes, which distinguishes it from other Eurasian marmots, which to varying degrees gravitate to mountainous and other types of dissected terrain. The species is described from the territory defined in the first description as “Poland” (Müller, 1776). Many researchers (Ognev, 1947, etc.) considered the place of the first description of the baibak to be the Right-Bank Ukraine, taking into account the geopolitical division at that time. The modern natural range of *M. bobak* is almost 3000 km long from west to east, and about 800 km from north to south. In this range today three subspecies are distinguished: nominative “European, or western”, “Kazakh, or eastern” (*M. b. schaganensis* Bashanov, 1930) and “Volga” (*M. b. kozlovi* Fokanov, 1966). The latter subspecies is not accepted by all specialists (Mashkin, 1997). The nominal subspecies is described from the territory west of the Dnieper, where it has been long extinct, the “Kazakh or eastern” subspecies from the southwestern part of the Orenburg region, and the Volga subspecies from the Saratov right bank. Thus, modern interpretations of their distribution form the following, rather paradoxical, picture. Modern populations of baibak, unanimously attributed to *M. b. bobak*, are separated from the supposed area of its original description by the Dnieper River. Its distribution is defined as the “European part of the range”, which in different interpretations reaches the Volga or includes the Volga region up to the Urals (Zarubin et al., 1996; Pavlinov and Lisovsky, 2012; Kryštufek and Vohralík, 2013). The Trans-Volga populations, which today are attributed to *M*.*b. bobak*, may actually belong to *M*.*b. shaganensis* - in any case, this subspecies was described from the Trans-Volga region. The range of *M*.*b. kozlovi* has been described both in the right bank of the Volga or Volga region without clearly defined boundaries (Gromov and Erbaeva, 1995; Zarubin et al., 1996; Mashkin, 1997) and eastward to the left bank of the Volga, despite the localization of its terra typica on the right bank (Kryštufek and Vohralík, 2013). Finally, the diversity of populations in the Trans-Ural, separated from the rest of the range by geographic-ecological barriers and unanimously attributed by all to *M*.*b. shaganensis*, has not been analyzed in detail, although the Trans-Urals part of the species’ range is extensive and may also be heterogeneous (Rumyantsev and Brandler, 2023). The presumably more complex subspecies composition of *M. bobak* is indicated by the identification of four “geographic populations” based on the analysis of the characteristics of the acoustic alarm calls of marmots from 11 points of the range (Nikol’skii, 2002).

Steppe marmot is an important commercial species on the one hand and an object of increased protection as a vulnerable species on the other. A thorough understanding of the structure of intraspecific variability and a well-founded subspecies taxonomy are important for planning rational use and conservation strategies. Despite close attention to the steppe marmot by various specialists, its genetic structure remains virtually unstudied. For the first time, we investigated the genetic variability of *M. bobak* using three mtDNA marker fragments throughout the species’ range, including material from terra typica subspecies.

## MATERIALS AND METHODS

### Tissue Sampling

Tissue samples of *M. bobak* were obtained from the large-scale research facility “Collection of Wildlife tissues for genetic research” of the Koltzov Institute of Developmental Biology, Russian Academy of Sciences (CWT IDB, state registration number 3579666). A significant part of the material was collected by the authors. Two *cytb* sequences (AF143916 and AF143917) of steppe marmots from our collection were obtained from the NCBI GenBank. We studied in total 110 individuals of *M. bobak* with recorded GPS coordinates of capture points distributed throughout the species’ range (Fig. 1; Supporting Information, Table S1). *COI, cytb* and *CR* sequences extracted from mitochondria deposited in GenBank with accession numbers OR085844 (*M. baibacina*) and OR085845 (*M. sibirica*) were used as an outgroup.

**Figure 1.**
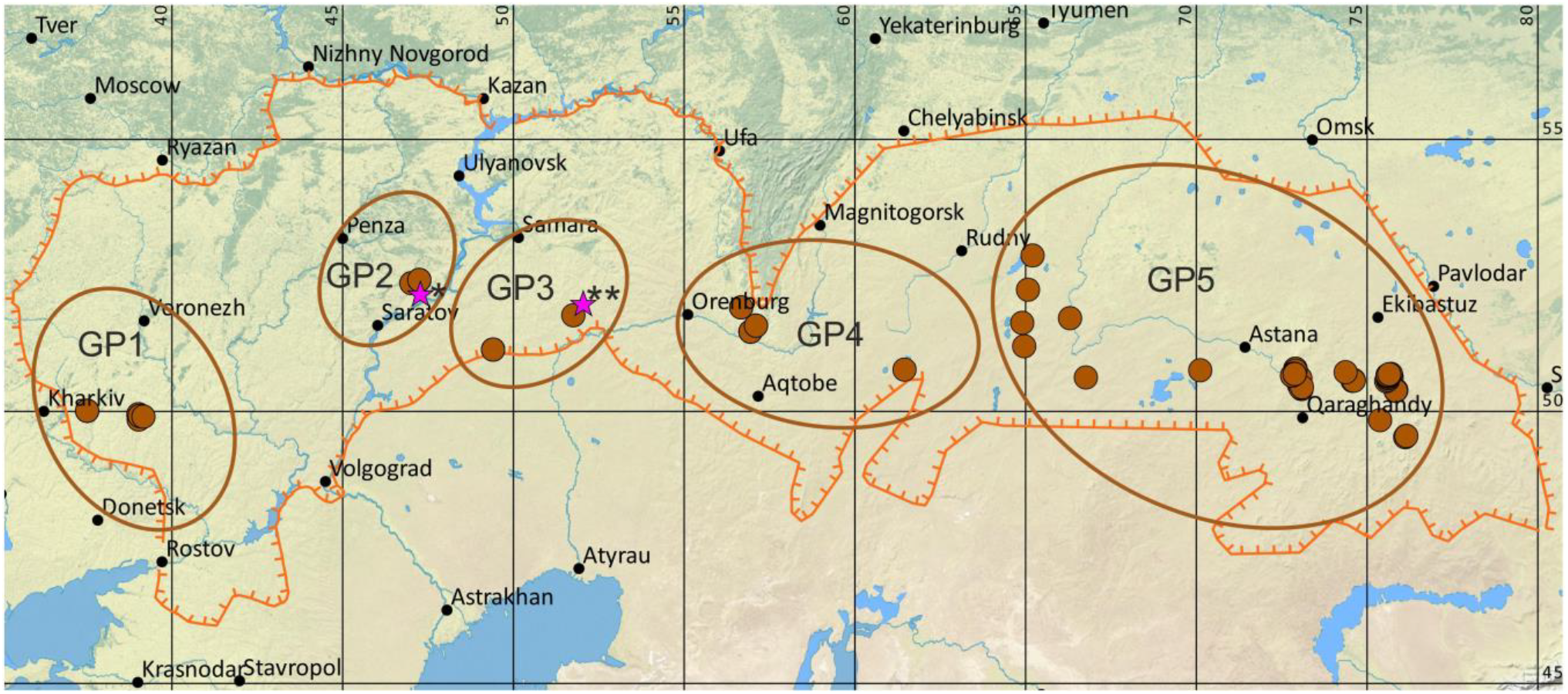
Distribution range (after Ognev 1947; Kryštufek et al. 2009 with our modifications) and localization of the studied populations of *M. bobak*. The orange line shows the range boundary; brown ovals and GP1-GP5 denote geographic populations (see text, and Table S1). Pink stars denote terra typica of subspecies: * *M*.*b. kozlovi*, ** *M*.*b. shaganensis*.

To analyze the influence of spatial and geographic factors on the structure of the species, we divided the entire sample into geographic populations (Naumov, 1971; Nikol’skii, 2002) (Fig. 1; Table S1). Local populations located on the territory isolated from other regions of the range by large physical geographic barriers were combined into geographic populations. Five geographic populations were identified: GP1 - Western, uniting populations of the Central Russian Upland; GP2 - Volga (populations of the Volga Upland right bank of the Volga River); GP3 - Transvolga (populations of the left bank of the Volga River and Obshchy Syrt); GP4 - South Ural (populations of the southern foothills of the Urals Mountains and Northern Mugodzhars up to the Turgay Depression in the east); GP5 - Kazakh (populations east of the Turgay Depression, including the Kazakh Uplands).

The animals were treated according to established international protocols, as in the Guidelines for Humane Endpoints for Animals Used in Biomedical Research. All the experimental protocols were approved by the Ethics Committee for Animal Research of the Koltzov Institute of Developmental Biology RAS in accordance with the Regulations for Laboratory Practice in the Russian Federation, the most recent protocol was No 37-25.06.2020. All efforts were made to minimize animal suffering.

### DNA Extraction, Amplification, and Sequencing

Genomic DNA was extracted by the standard salt method (Aljanabi & Martinez 1997). We used three mtDNA fragments as markers of intraspecific genetic variability: part of cytochrome oxidase subunit 1 (*COI*), full-length cytochrome *b* (*cytb*) and full-length control region (*CR*).

Amplification of each of the mtDNA marker sequences was performed using specific primers: SpCOXd and SpCOXr (Ermakov et al., 2015) for fragment *COI*; L14725 (Steppan et al., 1999) and H15915 (Harrison et al., 2003) for full-length *cytb*; MDL1 (Ermakov et al., 2002) and H00651 (Kocher et al., 1989) for full-length CR. In some cases, we used modified specific primers (for *cytb* or *CR*) with 5’ ends complemented by sequence primers (M13f and M13r, Ekimova et al., 2015). PCR was performed in 15 μl reaction mixture of Screen Mix or HS Taq DNA-polymerase (Evrogen, Russia) in a Veriti Thermal Cycler (Applied Biosystems, USA), at the following conditions: +95 °C – 20 sec, annealing temperature (specific to each pair of primers) – 40 sec, +72 °C – 40 sec (35 cycles), and final synthesis +72 °C – 10 min.

The sequencing reaction was performed in 10 μl of the reaction mixture using BigDye v.3.1 kit (Applied Biosystems, USA) according to the manufacturer’s protocols and external (M13f, M13r) or specific primers. The obtained fragments were analyzed in Formamide on an ABI 3500 automatic genetic analyzer (Applied Biosystems, USA). Laboratory procedures were performed on the basis of the Core Centrum of IDB RAS.

All newly sequenced haplotypes were deposited in GenBank with accession numbers *COI*: PQ670280-PQ670293, *cytb*: PQ657681-PQ657715, CR: PQ657716-PQ657760, (details see in the Supporting Information, Table S1).

### Molecular Data Analysis

The consensus sequences for *COI* (680 bp), *cytb* (1140 bp) and *CR* (995-996 bp) and concatenated sequences *COI*-*cytb*-*CR* (2815-2816 bp) were aligned with the algorithm MUSCLE implemented in the MEGA X software (Kumar et al. 2018).

Phylogenetic trees were inferred with concatenated *COI-cytb-CR* sequences under maximum likelihood (ML) and Bayesian criteria. The best model for nucleotide sequence evolution under BIC was selected separately for the 1st and 2nd codon positions of the coding part (part 1), the 3rd codon position of the coding part (part 2) and the non-coding part (part 3) of the concatenated sequence using ModelFinder (Kalyaanamoorthy et al., 2017). The following models were used to construct the trees: K3Pu+F+I (Kimura, 1981) for part 1, TN+F+G4 (Tamura and Nei, 1993) for part 2, HKY+F+I+G4 (Hasegawa, Kishino and Yano, 1985) for part 3. ML trees were reconstructed in the online version of IQ-TREE (http://iqtree.cibiv.univie.ac.at/) (Trifinopoulos et al., 2016). Bayesian tree reconstructions (MB) were inferred in MrBayes v3.2.6 (Ronquist et al., 2012) based on 2×10^6^ generations with each 5000th retained in two MCMC chains and discarding of the first 25%. Reconstruction was completed with a standard deviation of the separated frequencies of <0.01. Clade stability was tested using Ultrafast Bootstrap (Hoang et al., 2018) with 1000 replicates in ML trees and Bayesian posterior probabilities in MB trees. Bootstrap value of 70% and posterior probabilities of 0.9 were taken as the lowest significant value for supports of branches. Tree visualization was carried out in the FigTree v1.4.3 (http://tree.bio.ed.ac.uk/). Genetic differences were estimated by pairwise distances (p-distance, *pd*) in MEGA X.

Evolutionary network was constructed using the median joining (MJ) algorithm for *COI, cytb, CR* and concatenated *COI-cytb-CR* haplotypes using HaplowebMaker (https://eeg-ebe.github.io/HaplowebMaker/, accessed December 06, 2024) (Spöri et al., 2020).

The genetic variability indices including the number of haplotypes (*H*), the haplotype diversity (*Hd*) and the nucleotide diversity (*π*)] were estimated for *COI*-*cytb*-*CR* in DnaSP v.6.12.03 (Rozas et al., 2017). Tajima’s *D* and Fu’s *F*_*S*_ neutrality tests (Tajima, 1989, Fu, 1997) were also calculated to evaluate past population growth, decline or stability using DnaSP v.6.12.03. The expansion coefficient (*S/k*) was calculated to assess the differences between recent and historical population sizes, as the ratio of the number of variable sites (*S*) to the average number of pairwise nucleotide differences (*k*) (Peck & Congdon 2004).

## RESULTS

Analysis of 110 *COI*, 110 *cytb*, and 109 *CR* sequences revealed 14, 36, and 45 haplotypes among them, respectively (Supporting Information, Table S1).

Bayesian and ML phylogenetic trees of the combined *COI-cytb-CR* sequences had the same topology (Fig. 2). The separate *COI, cytb*, and *CR* trees had branching of major branches similar to the overall tree. The entire *M. bobak* sample was divided into two large clusters (phylogroups A and B in Fig. 2). Haplotypes of phylogroup A were found only in the eastern part of the range within the Kazakh geographic population (Fig. 3). Phylogroup B is subdivided into four phyletic lineages and its representatives are distributed throughout the entire range. The phyletic lineage B1 has a western distribution, from the western border of the range to the Obshchy Syrt. Haplotypes B2 genetically close to it are found together with haplotypes of B1 only in the westernmost populations, traditionally referred to the nominative subspecies of the steppe marmot. Sister phyletic lineages B3 and B4 are distributed consistently in the South Ural (B3) and Kazakh (B3 and B4) groups of populations in the central and eastern parts of the range (Fig. 3). Marmots captured in terra typica of subspecies *M*.*b. kozlovi* and *M*.*b. shaganensis* belong to the same phyletic lineage B1 (Figs. 2, 3).

**Figure 2.**
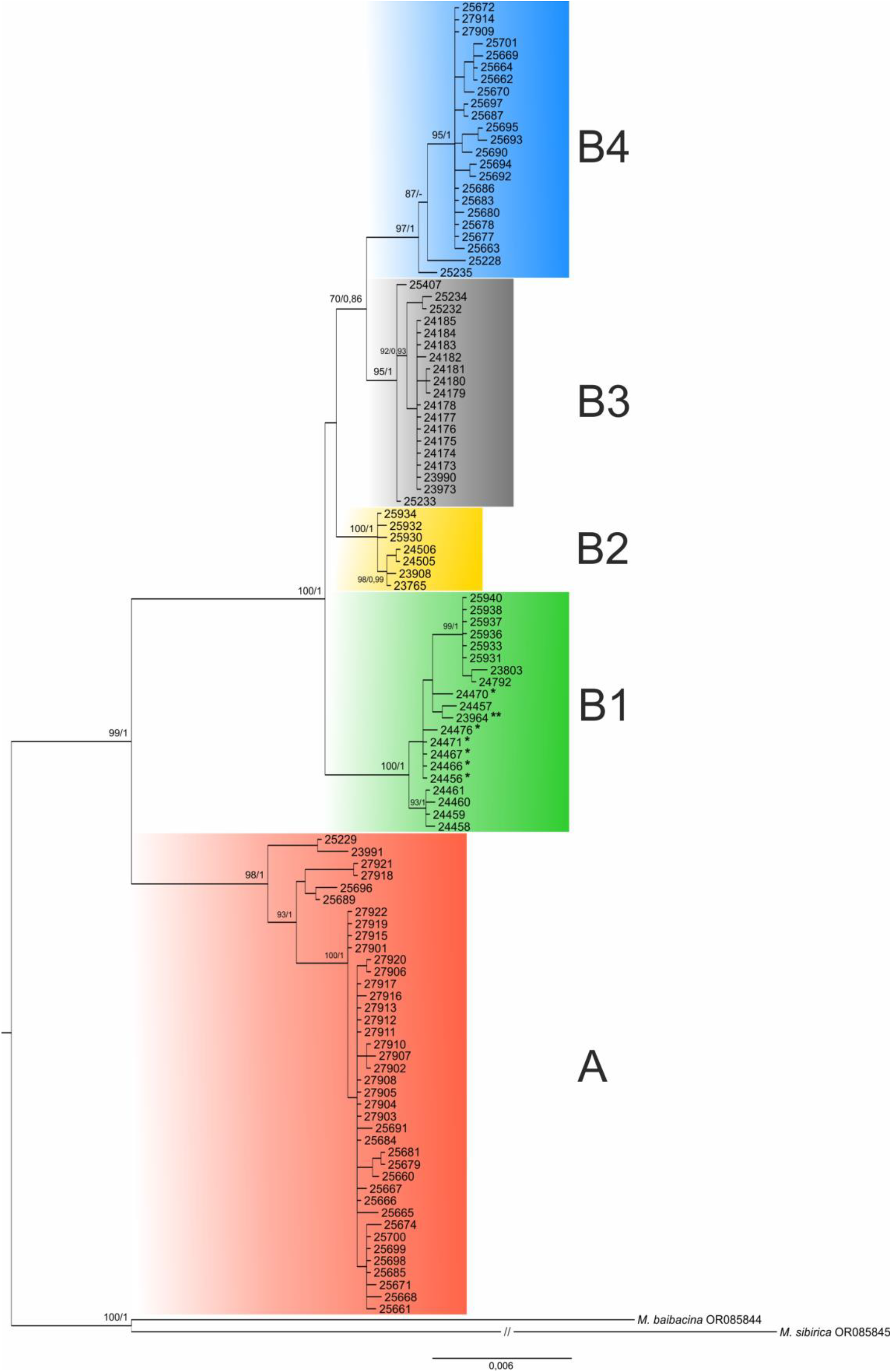
Phylogenetic tree of concatenated sequences *COI-cytb-CR* of *M. bobak*. Numbers above branches in tree represent UFboot support ≥70 of ML and posterior probabilities ≥0.9 of BIC. Labels A and B1-B4 denote phyletic lineages.

**Figure 3.**
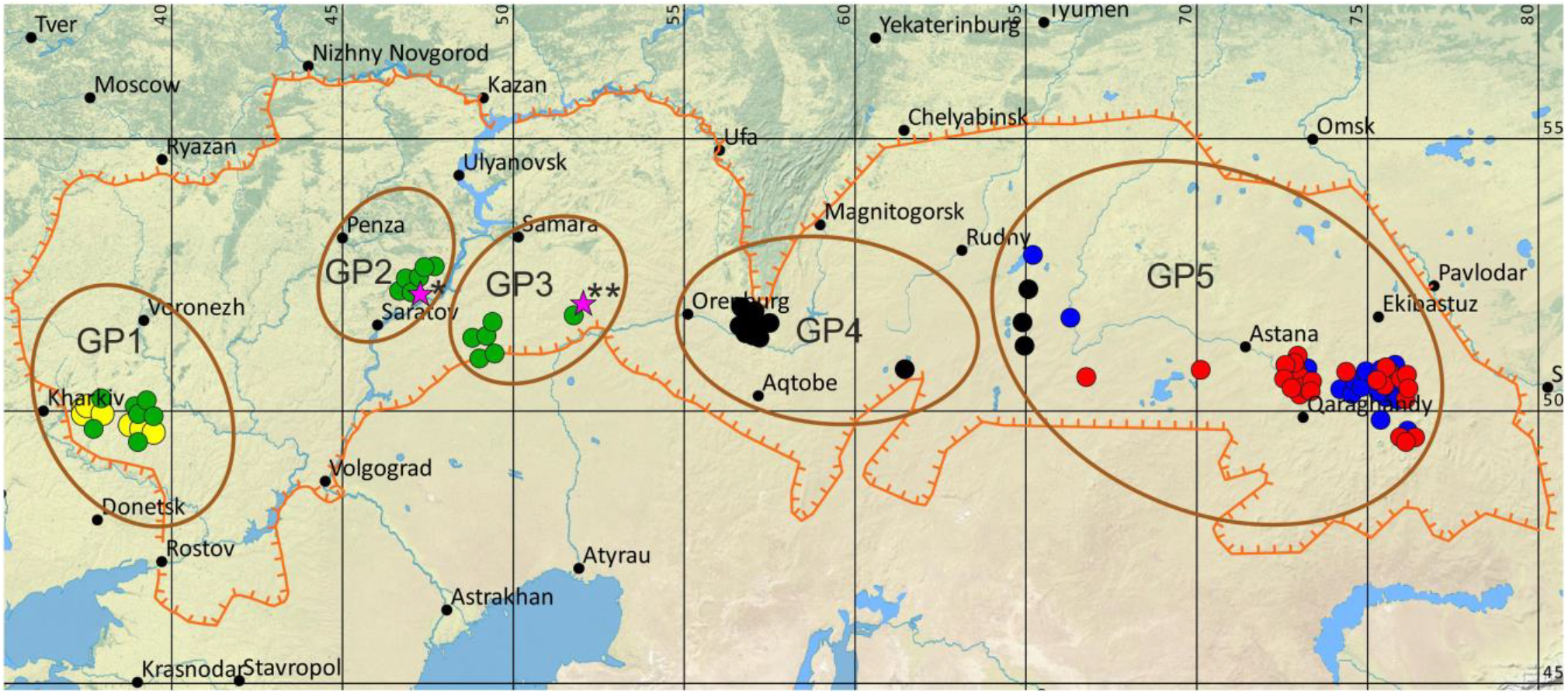
Map of localities studied for *M. bobak*. Colored dots indicate the localization of detected haplotypes (shown with an offset for clarity); colors correspond to Fig. 2. Other notations correspond to Fig. 1.

The evolutionary haplotype networks of individual markers and concatenated sequences (Fig. 4) generally correspond in the topology of phyletic lineages to the dendrogram (Fig. 2). The haplotypes of phylogroup A form a distant branch within each haplotype network, whereas the other phyletic lineages form several separate branches with few substitutions between them. The difference in the topologies of the networks consists of the different location of the haplotypes of lineages B1 and B4 on the *cytb* network (Fig. 4c) on one side and *COI* and *CR* (Fig. 4a, b) on the other relative to group A. This may be due to the lack of evolutionary intermediates or to the unevenness of substitutions in different parts of the mitochondrial genome. Apparently, such nonsynchrony determines low statistical support for the association of phylogroup B clusters (Fig. 2).

**Figure 4.**
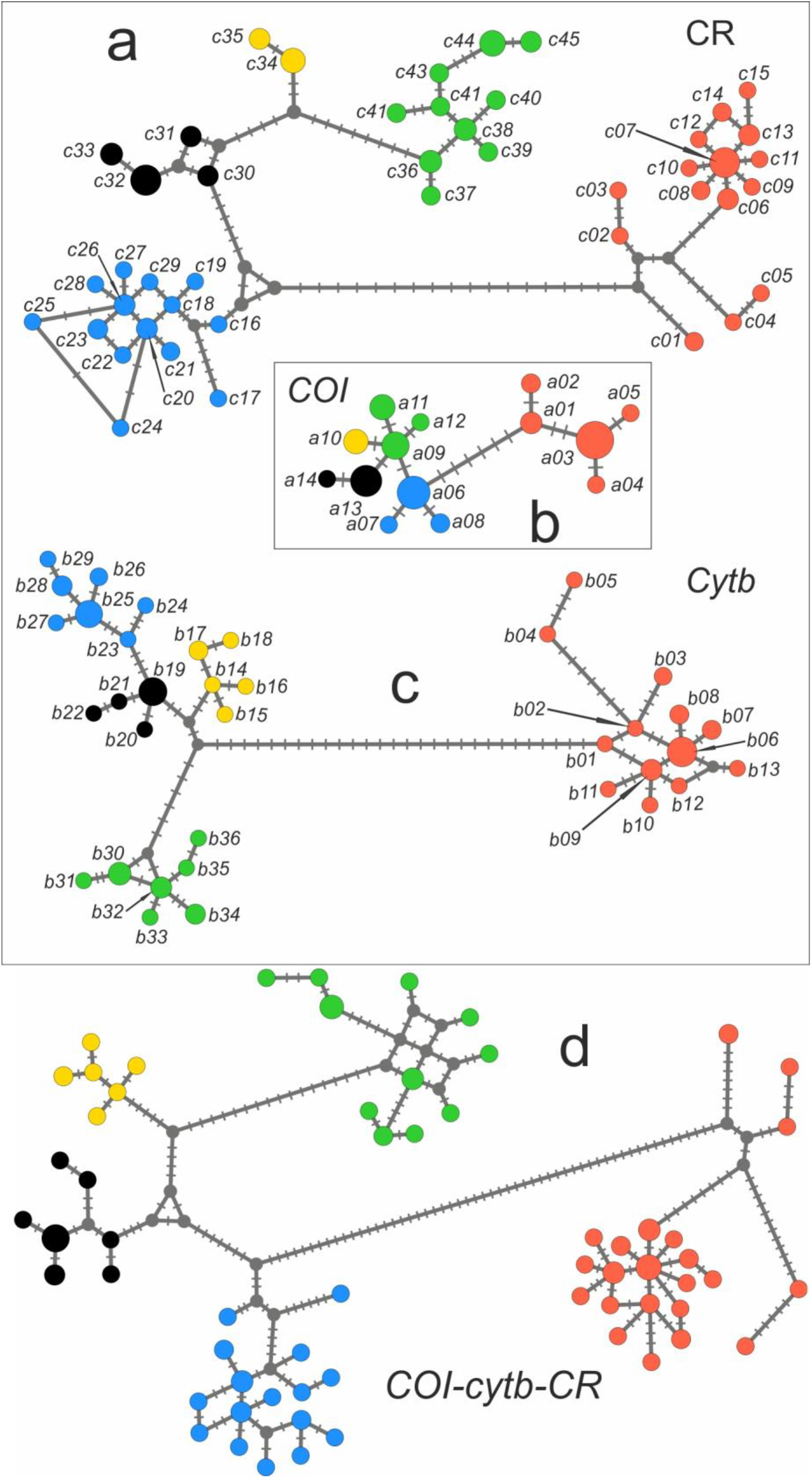
Evolutionary haplotype networks of (a) *CR*; (b) *COI*; (c) *cytb*; (d) concatenated sequences *COI-cytb-CR* of *M. bobak*. Haplotype labels are as in Table S1. Colors correspond to Figs. 2, 3.

The networks of all three markers, as well as the network of their concatenated sequences (Fig. 4d), show no complex connections between phylogroups, indicating their independent evolution, and the low number of substitutions between phyletic lineages B1-B4 suggests a recent time of their divergence.

Genetic differences of the selected geographical populations are presented in Table 1. The Kazakh population (GP5) differs from all other populations by an order of magnitude more than all other populations differ from each other. Also, intrapopulation differences in population GP5, which includes representatives of three phyletic lines (A, B3 and B4), are significantly higher than in the others. The genetic distances between phyletic groups A and B and between phyletic lineages B1-B4 with relatively low *pd* within clades (Table 2) indicate the established differentiated genetic structure of the species.

**Table 1.** Genetic differences of geographical populations (GP, see Supporting Information, Table S1) of *M. bobak*. Interpopulation genetic distances (*pd*) are shown below the diagonal, within-population *pd* are on the diagonal and standard error (S.E.) estimates are shown above the diagonal.

| G. P. | 1 | 2 | 3 | 4 | 5 |
| --- | --- | --- | --- | --- | --- |
| 1 | **0.0051** | 0.0010 | 0.0011 | 0.0014 | 0.0019 |
| 2 | 0.0054 | **0.0007** | 0.0006 | 0.0018 | 0.0021 |
| 3 | 0.0060 | 0.0017 | **0.0017** | 0.0018 | 0.0021 |
| 4 | 0.0079 | 0.0094 | 0.0097 | **0.0003** | 0.0020 |
| 5 | 0.0201 | 0.0206 | 0.0209 | 0.0184 | **0.0133** |

**Table 2.** Genetic distances (*pd*) between and within phyletic lines of *M. bobak*. Inter-group *pd* are shown below the diagonal, intra-group *pd* on the diagonal, standard error (S.E.) estimates are shown above the diagonal. * *pd* between A и B, ** *pd* within B. Phylogroup designations are as in Fig. 2.

|  | A | B1 | B2 | B3 | B4 | B total (±S.E.) |
| --- | --- | --- | --- | --- | --- | --- |
| A | **0.0024** | 0.0029 | 0.0029 | 0.0029 | 0.0027 | 0.0259±0.0026* |
| B1 | 0.0274 | **0.0021** | 0.0017 | 0.0018 | 0.0017 |  |
| B2 | 0.0247 | 0.0090 | **0.0007** | 0.0014 | 0.0017 |  |
| B3 | 0.0260 | 0.0096 | 0.0058 | **0.0005** | 0.0015 |  |
| B4 | 0.0247 | 0.0118 | 0.0089 | 0.0074 | **0.0015** | **0.0071±0.0009\*\*** |

Indicators of molecular diversity (Table 3) testified to diverse demographic processes in different parts of the range. High values of haplotypic diversity (*Hd*) combined with low nucleotide diversity (*π*) in the Western (GP1) and Trans-Volga (GP3) populations indicated their origin from a small ancestral population during a short time (Avise, 2000). The Volga (GP2) and South Ural (GP4) populations may have undergone long-term bottle-neck exposure. The high values of *Hd* and *π* in the Kazakh population (GP5) are obviously a consequence of the formation of its population from several previously isolated populations, which is consistent with the presence of individuals of several phyletic lineages in this area. The average values of the expansion coefficient in most populations indicated that the increase in their numbers occurred a relatively long time ago. The exception was population GP4, which appears to have experienced a recent population expansion. In general, range expansion or population growth was inferred for the entire sample based on the significance of *Fs* values and the non-significance of *F*\* and *D* values. A positive *Fs* value may indicate a recent bottleneck or enhanced selection for the species as a whole. At the same time, only population GP5 showed a recent expansion in *Fs*, while a positive significant *D* value in population GP1 may both indicate a sudden recent decline and reflect the presence of representatives of two phyletic lineages in the same population.

**Table 3.**
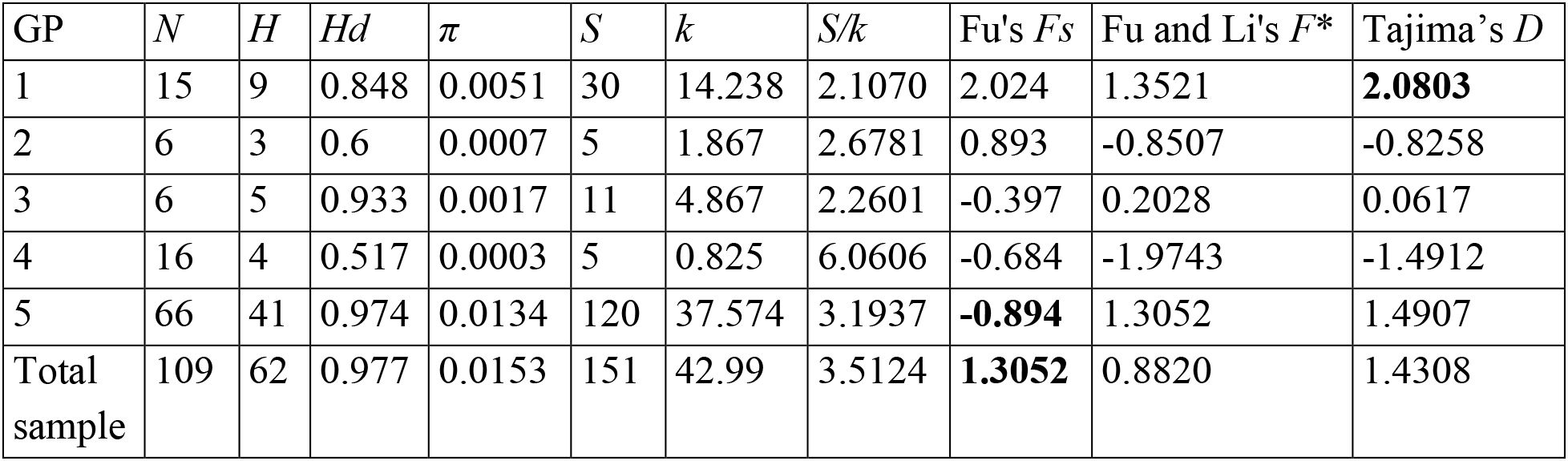
Summary of molecular diversity and neutrality test statistics indices by geographical populations (GP) and for the total sample of *M. bobak*: number of samples (*N*), number of haplotypes (*H*), the haplotype diversity (*Hd*), the nucleotide diversity (*π*), number of variable sites (*S*), average number of pairwise nucleotide differences (*k*), expansion coefficient (*S/k*), Fu and Li’s test and Tajima’s test results. Significant values are in bold.

| GP | *N* | *H* | *Hd* | $\pi$ | *S* | *k* | *S/k* | Fu's *F<sub>s</sub>* | Fu and Li's *F\** | Tajima's *D* |
| --- | --- | --- | --- | --- | --- | --- | --- | --- | --- | --- |
| 1 | 15 | 9 | 0.848 | 0.0051 | 30 | 14.238 | 2.1070 | 2.024 | 1.3521 | **2.0803** |
| 2 | 6 | 3 | 0.6 | 0.0007 | 5 | 1.867 | 2.6781 | 0.893 | -0.8507 | -0.8258 |
| 3 | 6 | 5 | 0.933 | 0.0017 | 11 | 4.867 | 2.2601 | -0.397 | 0.2028 | 0.0617 |
| 4 | 16 | 4 | 0.517 | 0.0003 | 5 | 0.825 | 6.0606 | -0.684 | -1.9743 | -1.4912 |
| 5 | 66 | 41 | 0.974 | 0.0134 | 120 | 37.574 | 3.1937 | **-0.894** | 1.3052 | 1.4907 |
| Total sample | 109 | 62 | 0.977 | 0.0153 | 151 | 42.99 | 3.5124 | **1.3052** | 0.8820 | 1.4308 |

## DISCUSSION

### Spatial genetic structure and history of *M. bobak*

In our study, we present the first range-wide assessment of genetic structure for the steppe marmot. Based on mitochondrial DNA data, our analyses revealed that the genetic structure of *M. bobak* consists of two main phylogroups. A rather high level of divergence between them suggests the presence of a long isolation between ancestral populations in the past.

The time of divergence between the *bobak* and *baibacina* lineages in the bobak group was dated to the Middle Pleistocene (1-0.5 Ma) (Mills et al., 2023). Fossil remains from the late Middle Pleistocene are known from the Caucasus, Crimea, right-bank Dnieper region (Gromov et al., 1965) and the Carpathians (Rădulescu and Kovács, 1970). Apparently, it is connected with the marmots’ exit to the plains from mountainous territories, which presumably were the primary landscape for marmots (Bibikov, 1989). Possibly, the original range of *M. bobak* was later divided in the meridional direction, because in the late Pleistocene up to the end of the Mikulin interglaciation (∼140-100 thousand BP), forests that reached the Caspian Sea prevailed in the Pre-Ural (Transvolga region) (Velichko, 2009; Svitoch, 2015), and the transition to the Würm glaciation was accompanied by the watering phase of the Turgay Spillway (Illarionov, 2013). Such isolation may explain the formation of the main phylogroups A and B in the gene pool of *M. bobak*. The expansion of marmots to the vast open spaces of eastern Europe and western Asia was associated with the formation of periglacial landscapes in the late Pleistocene, especially fully developed during the Valdai glaciation (Zimina and Gerasimov, 1980). The earliest mass fossil finds of *M. bobak* in the area were dated to this time (Rumyantsev and Markova, 2000). Our data indicated a wide distribution of phyletic lineages of group B both in the west and east of the range. This can be explained by the eastward expansion of steppe marmots from Eastern European populations, rather than westward expansion from southern and eastern Asian populations, as previously suggested (Zimina and Gerasimov, 1980).

The formation of phyletic lineages of phylogroup B probably occurred under the influence of short-term habitat fragmentation due to changing paleoclimatic conditions. Apparently, the B4 lineage was separated from the main trunk of the western steppe marmots by the last flooding of the Turgay Depression in the Late Karginian time ∼25 thousand years BP (Illarionov, 2013). Despite the drainage of the Turgay Spillway, the territory of the hollow in many places is still traversable for marmots only to the north of Lake Kushmurun (Kusmuryn) (Rumyantsev, 1991). This may explain the limited penetration of representatives of phyletic lineage B3 to the east of the Turgay Depression (Fig. 3).

The formation of the modern genetic structure of steppe marmot under the influence of reintroduction deserves separate consideration. By the middle of the 20th century, there was a multiple decrease in the number and range of the steppe marmot (Bibikov et al., 1990). To recover this species, large-scale reintroduction was carried out in the USSR in 1977-1993. A total of about 44800 marmots were dispersed to more than 415 locations (Rumyantsev et al., 2012). Such a large-scale artificial migration should have resulted in enhanced gene flow between distant populations and mixing of haplotypes of different phyletic lineages in the populations. Against expectations, our data did not support such a scenario. We analyzed information on marmot relocations at the sites of origin of the material we studied. The presence of two phyletic lineages B1 and B2 in GP1 populations (Fig. 3) could be the result of marmot migration from GP2 and/or GP3. However, we were unable to find evidence of such translocations. According to literature data, no translocations were made to western populations (Dezhkin et al., 1983), and all cases of marmot translocations in these regions were limited to either intra-regional movements or reintroduction from the studied populations to western regions (Tokarsky et al., 2006). Movements of marmots from the Volga populations (GP2) to the Trans-Volga territories occurred within the coastal areas. The populations of the eastern districts of the Saratov Region (Ozinsky District), 150 km away from them, from which the material we studied comes, were always considered relict and marmots were not introduced into them (Semikhatova and Khrustov, 1996; Kondratenkov et al., 2020). In the Orenburg region, 1451 marmots were settled in eight districts in 1981-1986 (Rudy, 1988), and 1883 individuals in 1991-1996 (Fedorenko, 2009). The donors were populations from the eastern districts of the region (GP4). Relocation was carried out to the central and western parts of the region. Traces of this reintroduction did not appear in our material, probably due to the limited sample. At the same time, the success rate of these reintroductions was very low (Rudy, 1988), with marmots’ survival rate of 10% (Fedorenko, 2009). This may be why they have not contributed to the diversity of the species gene pool. In Kazakhstan, most relocations were either intra-regional or between neighboring regions (Bauer E.L., Neldner V.V., personal communication) and could not significantly change the spatial and genetic structure of populations in the eastern Kazakhstan part of the range.

### Taxonomic implications

The results obtained did not agree with the accepted subspecies systematics of *M. bobak*. Marmots from western populations, traditionally belonging to the nominative subspecies, as well as individuals from *terra tipica* of the other two described subspecies *M*.*b. kozlovi* and *M*.*b. shaganensis* have mtDNA haplotypes belonging to the same phyletic lineage B1 (Figs. 2, 3). In addition, B2 haplotypes are present in the western populations. At the same time, all other genetically differentiated forms of steppe marmots belonging to other phyletic lineages cannot be formally attributed to any of the described subspecies of *M. bobak*. At the same time, haplotypes of only one phyletic lineage B3 were found in the populations of the Southern Urals (GP4), while marmots inhabiting the territory east of the Turgay Depression (GP5) are genetically heterogeneous and peculiar. The presence of individuals with three different types of mitochondrial genome in the latter area indicates the probable mixing of distant evolutionary lineages and increased heterozygosity of the marmot nuclear genome in these populations. We believe that, contrary to prevailing ideas, all marmots from GP4 and GP5 cannot belong to the Kazakh subspecies *M*.*b. shaganensis*, with individuals from *terra tipica* of which they are clearly genetically differentiated. Our results do not support the separation of subspecies *M*.*b. kozlovi*. The genetic uncertainty of the subspecies structure of the species is consistent with the uncertainty of morphological subspecies diagnoses. The delineation of subspecies based on traits with clinal variability and the lack of qualitative differentiating characters forced the authors of a recent revision of ground squirrels to abandon the division of steppe marmots into subspecies (Kryštufek and Vohralík, 2013). The problem of subspecies definition of marmots of the eastern part of the range and the boundaries of nominative subspecies requires additional research and thorough revision.

## CONCLUSION

Our study revealed genetic intraspecific heterogeneity in the steppe marmot for the first time. The revealed genetic structure of the species turned out to be non-obvious and allows assuming large-scale expansions both in the western and eastern directions within the historical range of *M. bobak* in the past. Also, the character of intraspecific genetic variability is not consistent with the accepted subspecies division. Additional genetic, morphological, ecological and other studies are required to create an adequate subspecies system of the steppe marmot.

